# Phototaxis as a Collective Phenomenon in Cyanobacterial Colonies

**DOI:** 10.1101/155622

**Authors:** P. Varuni, Shakti N. Menon, Gautam I. Menon

## Abstract

Cyanobacteria are a widely distributed, diverse group of photosynthetic bacteria that exhibit phototaxis, or motion in response to light. Cyanobacteria such as *Synechocystis* sp. secrete a mixture of complex polysaccharides that facilitate cell motion, while their type 4 pili allow them to physically attach to each other. Even though cells can respond individually to light, colonies of such bacteria are observed to move collectively towards the light source in dense finger-like projections. Agent-based models are especially useful in connecting individual cell behaviour with the emergent collective phenomena that arise out of their interactions. We present an agent-based model for cyanobacterial phototaxis that accounts for slime deposition as well as for direct physical links between bacteria, mediated through their type 4 pili. We reproduce the experimentally observed aggregation of cells at the colony boundary as a precursor to finger formation. Our model also describes the changes in colony morphology that occur when the location of the light source is abruptly changed. We find that the overall motion of cells toward light remains relatively unimpaired even if a fraction of them do not sense light, allowing heterogeneous populations to continue to mount a robust collective response to stimuli. Our work suggests that in addition to bio-chemical signalling via diffusible molecules in the context of bacterial quorum-sensing, short-ranged physical interactions may also contribute to collective effects in bacterial motility.

## Introduction

A complex set of sensory and regulatory pathways drive decision-making by micro-organisms. For motile micro-organisms, such processes can result in an overall motion towards or away from a host of stimuli. The most well-examined among these behaviours is chemotaxis, studied extensively in flagellated *Escherichia coli* which swim up (or down) chemical gradients [1]. While chemotaxis is relatively well understood, the mechanisms by which various micro-organisms respond similarly to many other types of stimuli [2] including pH changes [3], oxygen [4], osmolarity [5], light [6] and magnetic fields [7] are an area of active research.

Phototaxis, or motion in response to a light stimulus, was first reported over a century ago in eukaryotic photoautotrophs [8–11]. Recent studies on this phenomenon have focused on cyanobacteria or ‘blue-green algae’, which are a widely distributed, diverse group of oxygenic photosynthetic gram-negative bacteria. The model cyanobacterium *Synechocystis* sp. PCC 6803 displays robust positive phototaxis in which dense finger-like projections of cells emanate from a colony over a period of 1-3 days, and move toward a source of white light. Specific wavelengths of light elicit responses that range from slower moving colony fronts for red and far-red light [12] to negative phototaxis under blue, UV and high light conditions [13]. A wide range of wavelength and intensity-dependent tactic responses to light stimuli have been observed in other cyanobacterial species [12, 14].

A specific set of genes, involved in the production and extrusion of complex polysaccharides (‘slime’), are essential for *Synechocystis* motility [15]. *Synechocystis* possess multifunctional type 4 pili (T4P) that allow them to attach to substrates as well as other cells. These bacteria exhibit “twitching” or “gliding” motility, involving slime secretion. Gliding motility is slow, with speeds ranging from 0.03 to 0.07 *μ*m/s [16]. Such speeds are far slower than typical flagella-mediated motion which occurs at speeds of 20 to 50 *μ*m/s [17]. Phototaxis in *Synechocystis* colonies occurs in two distinct phases. Initially, individual cells move toward the edge of the colony closest to the light source, forming a crescent of cells. In a subsequent step, cells move towards the light source in regular, dense finger-like projections (see Fig. 1 of [18]). Studies that track the motion of individual *Synechocystis* cells following the application of a directional light source have shown that such cells initially move towards the light source individually [19]. Subsequently, their motion becomes density-dependent [16]. Cell motion at early times is similar to a random walk motion biased in the direction of the light source. This bias increases as cells aggregate into smaller motile groups, eventually leading to the formation of finger-like projections in which the directional bias is most pronounced. When these fingers intersect with the path of a previously formed finger, the cell speed increases, likely a result of encountering the slime that normally accompanies T4P-mediated motility. That even small aggregations of cells (5-8) exhibit an increased bias in the direction of the light source [16] suggests that the “social” aspect to phototaxis might be mediated by physical connections between cells. Similar social phenomena have been documented in other T4P systems such as *Myxococcus xanthus* [20, 21].

**Figure 1:**
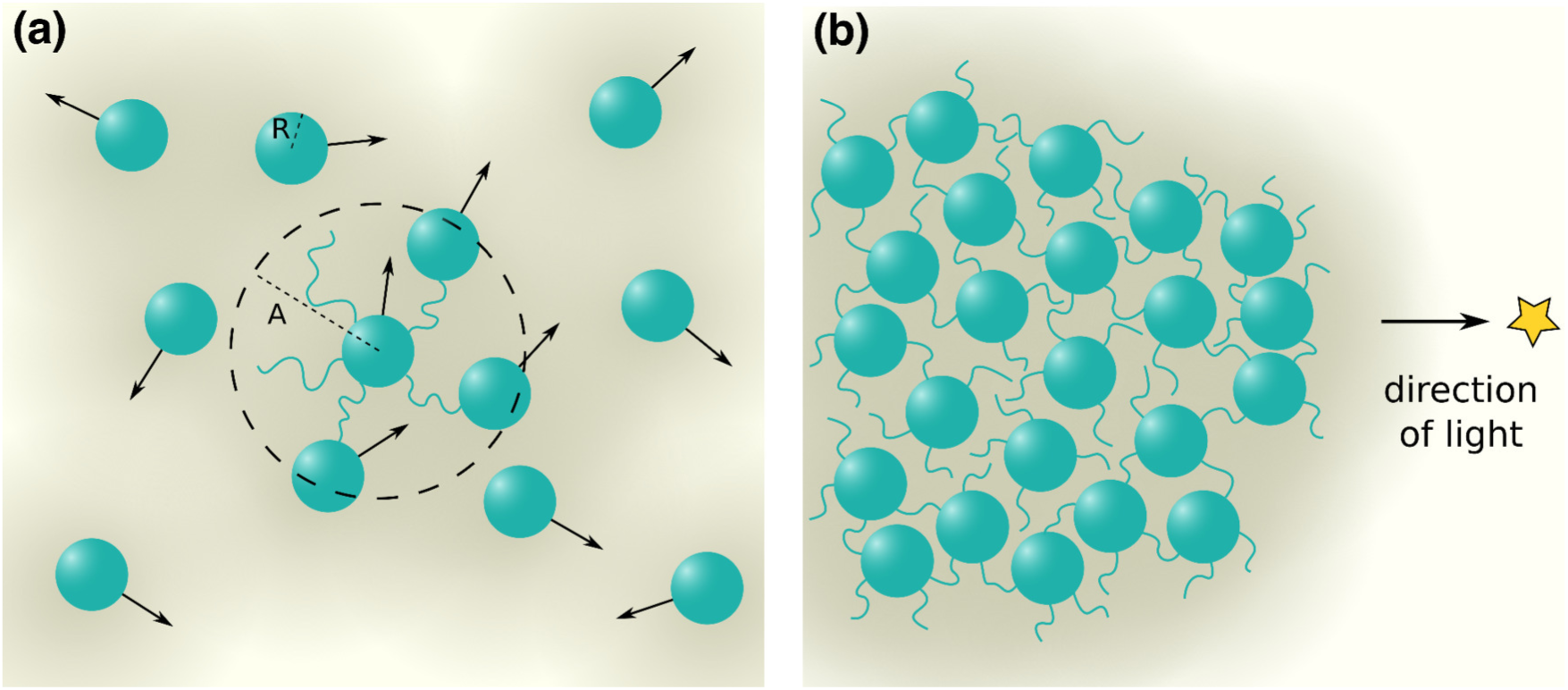
Schematic of simulation system. (a) Cells are represented as green spheres of radius *R*. The amount of slime is proportional to the intensity of background colour (darker implying more slime). Cells can attach to other cells through TFP, which can extend to length *A*. (b) In the presence of a distant light source (indicated by a star), the colony morphology gradually changes.

Recent mathematical models of phototaxis assume that cells move via a random walk biased in the direction of a light source. In order to investigate the role that slime plays in phototaxis, these models assume that cells prefer to move on regions of the substrate that have already been traversed by other cells, based on observations that fingers break up on entering an area with pre-existing slime [16]. This framework has been extended to include density-dependent interactions between neighbouring cells [16, 22]. It was shown in a cellular automaton model that cell aggregates move collectively towards the light source in finger-like projections, a result verified using a stochastic model with similar rules [23]. Other models emphasize the role of physical attachments between cells [22, 24, 25]. These models show that allowing frequent detachment and reattachment to neighbouring cells leads to increased aggregation. Varying the range across which cells interact modulates the dynamics of taxis patterns. A recent approach uses a reaction-diffusion model to obtain finger-like projections from the cell colonies extended in the direction of light [26]. Here, slime was modelled in terms of a variation of surface properties, influenced by local cell concentrations.

While these models help to elucidate aspects of the collective motion of cyanobacterial cell colonies in response to light, several important questions remain. For instance, the physical interactions between cells appear to be significant in both the initial stages of aggregation as well as in finger formation. Furthermore, previous agent-based models of phototaxis typically assume cells to be point-like particles, thus preventing an investigation into the effects of density and crowding. A natural background for investigating the collective dynamics of phototaxis is is provided by the theoretical framework of active matter systems. Ever since the seminal model of Vicsek *et al.* [27], there have been numerous attempts to describe the motion of large aggregations of self-propelled particles, i.e. units whose movement is driven by an internal energy source [28–30]. This framework has been applied to the study of flocking dynamics, although its simplicity allows it to be utilized across a wide range of systems [31]. With regards to the collective motion of cells arising through taxis, most active matter descriptions have been limited to the context of bacterial swimming. These encompass both run and tumble [32] and active Brownian [33] models: two classes of active particle systems that exhibit similar macroscopic dynamics [34, 35]. However, we know of no comparable description in the context of phototaxis.

In this paper, we present an agent-based model for the collective motion of a cyanobacterial colony in the presence of a light source. As detailed in the Methods section, we describe the movement of individual cells, modelled as particles of finite extent, that are initially located within a slime-filled colony. We explicitly consider the role of T4P, as well as of slime deposition, on the resulting dynamics. We assume that cells move randomly with a bias in the direction of a light source, governed by a fixed probability. We investigate how variations in this probability affect the collective dynamics. In addition, we study how the colony behaviour changes upon increasing the fraction of cells that are unable to sense the light source. Such cells effectively act as “freeloaders” that can only move in a directed manner by latching onto cells that can sense light. Our model captures a number of reported observations of cyanobacterial colony behaviour. In addition, its flexibility implies that it can be used to provide predictions for several experimental contexts that have not yet been probed systematically.

## Methods

Our model simulates a colony of cells, each of which are capable of motion, that reside on a flat substrate. The simulation proceeds by describing how the positions of all cells are to be updated at successive time steps. The motion of cells is biased in the direction of a light source, if it is present, while they move randomly in its absence. As cells move, they secrete slime. The presence of slime reduces the friction encountered by cells as they move across the substrate, thus facilitating the motion of other cells across that region. We also assume that cells experience forces from other cells in their vicinity. These inter-cellular forces account for the collective aspects of phototactic motion.

### The cell

We describe each cell as a disc of radius *R*. Each cell *i* is specified by a two dimensional vector, **X**_*i*_, described through the coordinates (*x*_*i*_*, y*_*i*_). At every time step *t*, each cell can either move via phototaxis in the direction, 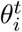, of an external light source with probability *p*_photo_, or in a random direction in the interval [0, 2*π*] with probability 1 *-p*_photo_. Time steps are separated by Δ*t*, which we set to 1. The maximum distance that a cell in a slime-rich background can move in a single time step is a tenth of its radius. In slime-poor backgrounds, the reduced mobility of the cell implies that it moves a smaller distance in the same time.

### Slime Deposition

We model the secretion of slime by considering a regular square lattice that underlies the cells. The lattice point at row *r* and column *c* is specified by (*r, c*). At each time step, every cell deposits slime. The slime content *S*^*t*^ of the lattice point closest to the cell’s centre is thus incremented by an amount *S*_*rate*_:

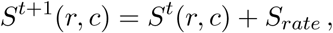

where *S*_*rate*_ is the rate of deposition of slime. *S*^*t*^(*r, c*) can increase up to *S*_*max*_, the maximum amount of slime that a grid point can contain. We assume that (i) slime once deposited at a lattice point remains there permanently, i.e. it does not decay, and (ii) slime does not diffuse to neighbouring lattice points. This latter is a valid assumption for a dense gel, given the time scales over which our simulation proceeds. Furthermore this assumption allows, in principle, for the creation of steep gradients in slime content.

### Cell-cell interactions

The presence of neighbouring cells modulates the direction of motion of a cell. Each cell has a fixed number *a* of T4P. These pili can attach to other cells lying within a certain distance *A* of the cell edge (see Fig. 1). We assume that these T4P links are temporary - they break and re-form at each new time step of the simulation - and that each cell can have at most *a* links with other cells. For the duration that a cell pair (*i, j*) remains attached, cell *j* exerts a force **f**_*ji*_ on *i* with magnitude *K*_*ij*_ that depends on the distance between cell *i* and *j*, *D*_*ij*_, in the following way:

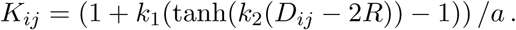

In order to discourage overlaps between cells, we use a sigmoidal form for *K*_*ij*_ that is negative at short distances, implying a repulsive force. The magnitude of the force is determined by the parameter *k*_1_, and the division by the parameter *a* accounts for the fact that each cell distributes its energy across their *a* pili to exert forces. The parameter *k*_2_ controls the slope of the sigmoidal function, and hence determines how sharply the magnitude of force reduces as inter-cell distance increases. Note that when *D*_*ij*_ *<* 2*R*, i.e. the cells overlap, this functional form results in a repulsive force, which is to be expected. In other words, the force term incorporates soft-core repulsion between cells. The values of *k*_1_ and *k*_2_ were chosen by scanning through the parameter space for finger-like projections (see Fig. S1).

The cell experiences an external force, **F**_*i*_ = Σ_*j*_ *K*_*ij*_ **f_ij_** from other cells *j* in its neighbourhood. In addition, the tendency of the cell to move in the direction 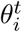 is modelled through an additional force 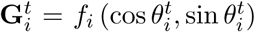, where *f*_*i*_ = 1. Thus, the force acting on a cell at each time step *t* is **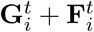**.

### Cell movement

As cell motion is facilitated by the presence of slime, we assume that the motility of a cell at any instant in time depends on the amount of slime at the grid point closest to the cell centre. The position of each cell is updated at the end of each time step through the equation of motion. Hence, the net result of external forces on a cell *i*, over a time interval Δ*t* is a change in the position of the cell, computed through the expression:

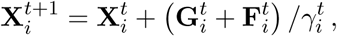

where 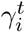 is a friction factor that is associated with the presence of slime lying below a cell. We assume that:

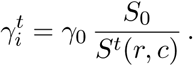

Here, *S*_0_ is the initial slime concentration within the colony while *S*^*t*^(*r, c*) is the slime content associated with site (*r, c*) at time *t*.

### Simulation details

We simulate the dynamics of a cyanobacterial colony containing 500 cells. The cells are initially distributed randomly in space over the extent of a circular or rectangular domain. We assume that the colony has an initially uniform slime distribution, with the slime content set at *S*_0_ for every grid point contained within the domain. We specify the initial positions (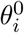) of each cell, their initial speeds, 1*/γ*_0_, and the angles 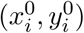 that they would move in if other cells were absent. Unless otherwise indicated, the parameters used are those listed in

Table 1.

**Table 1:**
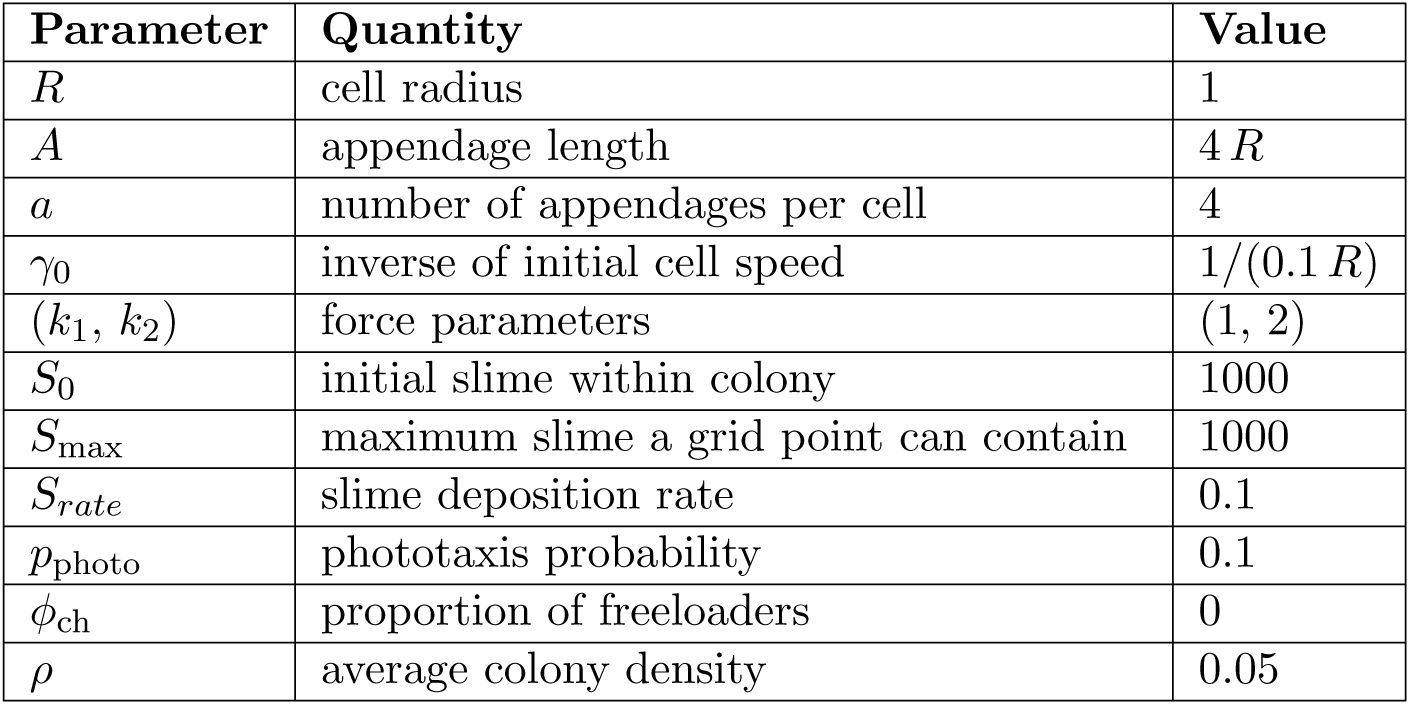
Parameters used in simulations (unless mentioned otherwise)

The length parameters used in these simulations were scaled to the size of a *Synechocystis* cell. These cells are about 1*μm* in radius [36], which is assumed to be one length unit in these simulations. Observations of cells under a SEM suggest that the T4Ps can be about four times times the cell radius and that they number about four [37]. The time parameters were scaled to the reported cell speed [38]. We have assumed that the highest cell speeds are obtained under maximum slime conditions. To our knowledge, there have not been any quantitative measurements of slime deposition or a detailed explanation of the mechanism through which slime affects cell speeds. Also, barring a few studies (e.g. [39]) the forces that cells can apply on each other through T4P have not been systematically measured. Hence, our choice of values for the associated parameters (*k*_1_*, k*_2_) was based on an exploration of the parameter space (see Fig. S1).

At each iteration of our simulation, we begin by updating the angles 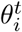 of all cells *i*, based on the probability of moving in the direction of light, *p*_photo_. We then find the set of all pairs of cells that are within tugging distance and determine which “tugs” occur. Next, we compute the distances between the *x* and *y* projections of all cells to determine the total force exerted by neighbouring cells on each other. The position of each cell is then updated using the equations of motion. At the end of each iteration, cells secrete a unit of slime in the grid point closest to their centre, and we update the slime matrix accordingly.

## Results

In Fig. 2, the left column displays the positions of cells in an initially circular colony at different times after the application of light, as indicated in the figure caption. The right column shows the trajectories of an individual cell, shown in purple against the background of trajectories of all other cells, shown in grey.

**Figure 2:**
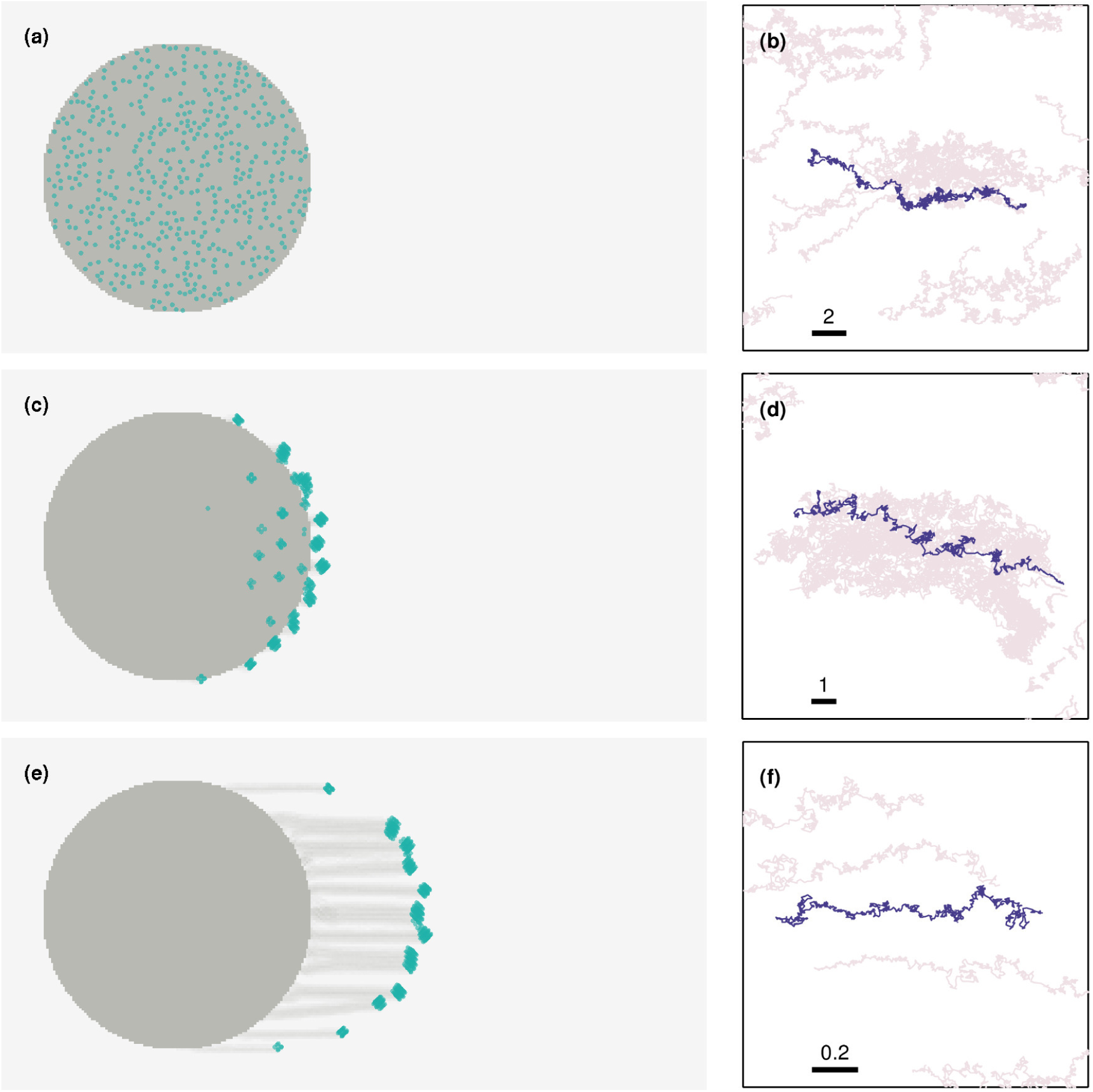
Finger formation and individual cell paths at different stages. The left column displays the position of each cell, indicated by green markers, while the concentration of slime is proportional to the intensity of colour (darker implying more slime). (a) At *t* = 0, the cells lie within a circular colony. In the presence of a light source placed at infinity (to the right of the colony), the colony morphology gradually changes. (c) At around *t* = 8 × 10^3^ finger like projections begin to form. (e) As seen from this snapshot at *t* = 50 × 10^3^, smaller projections can merge over time to form larger, well-defined projections. The right column displays the path of a representative cell (purple) and those of surrounding cells (gray) over an interval of 10^3^ steps. The thick lines represent the spatial scales of each of the panels (which is identical in both *x* and *y* directions), and their extent is denoted by the corresponding number on top. The cases shown are for (b) the initial trajectory (*t* = 0–10^3^), (d) early finger formation (*t* = 8 × 10^3^–9 × 10^3^), and (f) late finger formation (*t* = 5 × 10^4^–5.1 × 10^4^).

Fig. 2(a) shows the positions of individual cells at a time just after the application of light from a source at the right of the colony. Fig. 2(b) shows the trajectory of a labelled cell for 10^3^ time steps starting from this initial time. The overall bias of this trajectory towards the light source is obvious, although cells in the bulk feel forces that are largely isotropic from the cells in their vicinity. As Fig. 2(c) shows, cells concentrate into denser circular regions at later times. These cells move collectively towards the light source. The trajectory shown in Fig. 2(d) is now largely straight and directed toward the source. At a still later time, shown in Fig. 2(e), these dense accumulations of cells split off from the main colony although they remain connected to it through a trail of slime. A few isolated cells may remain in the bulk of the finger. The trajectory in Fig. 2(f) illustrates the far more directed motion of a cell within the finger-like protrusion, since it is now confined to a narrow strip of slime (See Movie S1).

Fig. 3 depicts properties of the trajectories of individual cells, initiated from a semi-infinite aggregate out of which cells move perpendicular to the surface upon application of light. Fig. 3(a) provides a snapshot of a configuration of an initially flat colony of cells moving in response to a light source at infinity, placed to the right of the colony. This is shown for a value of *p*_photo_ = 0.1, with a snapshot at *t* = 10^5^ time steps. Fig. 3(b) shows a rose plot of the direction of motion of individual cells obtained in the following way: each cell is tracked across a moving window and the net direction of displacement over five time steps is calculated. The angle this makes with respect to the x-axis is histogrammed and plotted. As can be seen, the anisotropy of the rose plot is indicative of the anisotropy of cellular motion induced by phototaxis towards the light source.

**Figure 3:**
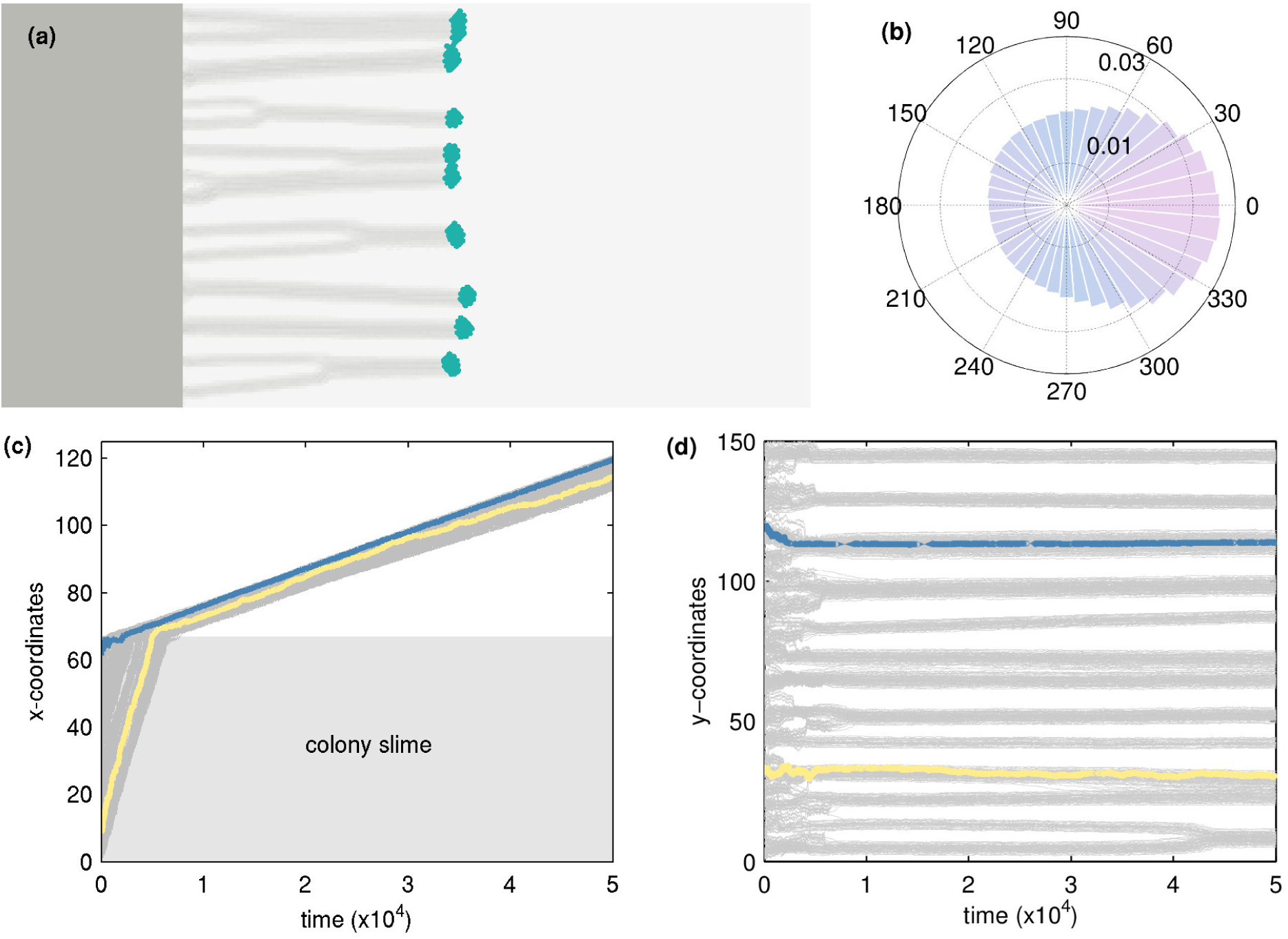
Phototaxis in the direction of light. (a) Snapshot of a flat colony of cells that move towards a light source placed at infinity (to the right of the colony) corresponding to a value of *p*_photo_ = 0.1 at *t* = 10^5^. (b) Corresponding rose plots for the net directions of the cells, (calculated over 5 time steps). (c-d) Trajectories of individual cells over time, with two randomly chosen cells coloured distinctly to illustrate characteristic paths. (c) The *x*-component of the trajectories, with the edge of the colony indicated by a dashed line. (d) The *y*-component of the trajectories, showing finger formation.

Figs. 3(c-d) depict the kymographs of the trajectories of individual cells. The cells at the boundary of the colony initially move slower than those in the bulk due to the lack of slime in their proximity. As the fingers form and cells in the bulk move towards the surface their velocities slow to match the cells at the boundary. We see substantial accumulation of cells at the boundary before fingers form and extend out of the colony, an indication of the importance of collective effects triggering finger formation. Fig. 4 shows the evolution of fingers from an initial flat colony when the probability of moving towards light changes across nearly two orders of magnitude. Surprisingly, even a relatively small bias (Fig. 4(a), *p*_photo_ = 0.01) produces fingers, although these tend to appear somewhat more disordered than fingers obtained at higher bias (Fig. 4(c), *p*_photo_ = 0.10 and Fig. 4(e), *p*_photo_ = 0.50). A larger *p*_photo_ leads to longer, better-defined fingers with a large cell density at the tip. Fig. 4(b,d,f) show the corresponding rose plots of the direction in which cells move. As seen in Fig. 4(f), at large *p*_photo_, cells mostly move in the direction of the light alone, leading to far more elliptical rose plots compared to the case of small *p*_photo_. These results suggest that very small *p*_photo_ can lead to robust phototaxis.

**Figure 4:**
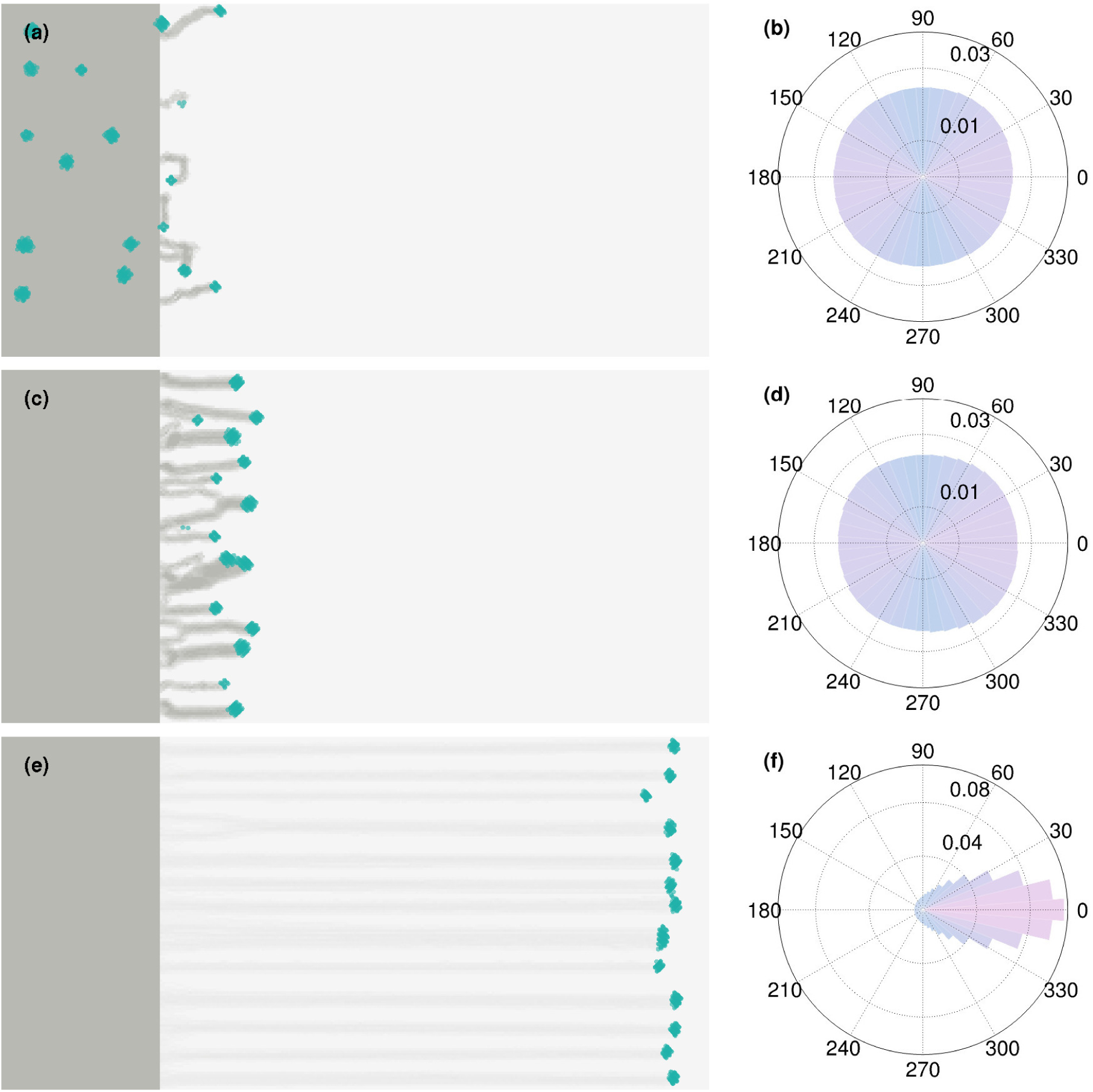
Robust phototaxis can occur even with a small bias for moving in the direction of light. The left column displays snapshots of flat colonies at *t* = 10^5^ for three different choices of *p*_photo_, namely (a) *p*_photo_ = 0, (c) *p*_photo_ = 0.01 and (e) *p*_photo_ = 0.5. We see that fingers can form even for very low *p*_photo_. The right column displays the corresponding rose plots for the net directions of the cells, (calculated over 5 time steps) showing that directed motion is enhanced at higher *p*_photo_.

It has been observed that cells increase their speed when they encounter regions in which slime has already been laid down. To simulate this, we initiate fingers from a colony allowing them to grow in the direction of light placed towards the right of the colony. After the fingers grow to a certain length, they encounter a band of slime placed normal to the direction of their growth. As seen in Fig. 5(a), after crossing the slime finger growth continues for those fingers that manage to reach the band. In Fig. 5(b) we show kymographs of the trajectories themselves. These show that cells that encounter the band of slime speed up within it before emerging and then move with the same velocity that they had before encountering the slime band (See Movie S2).

**Figure 5:**
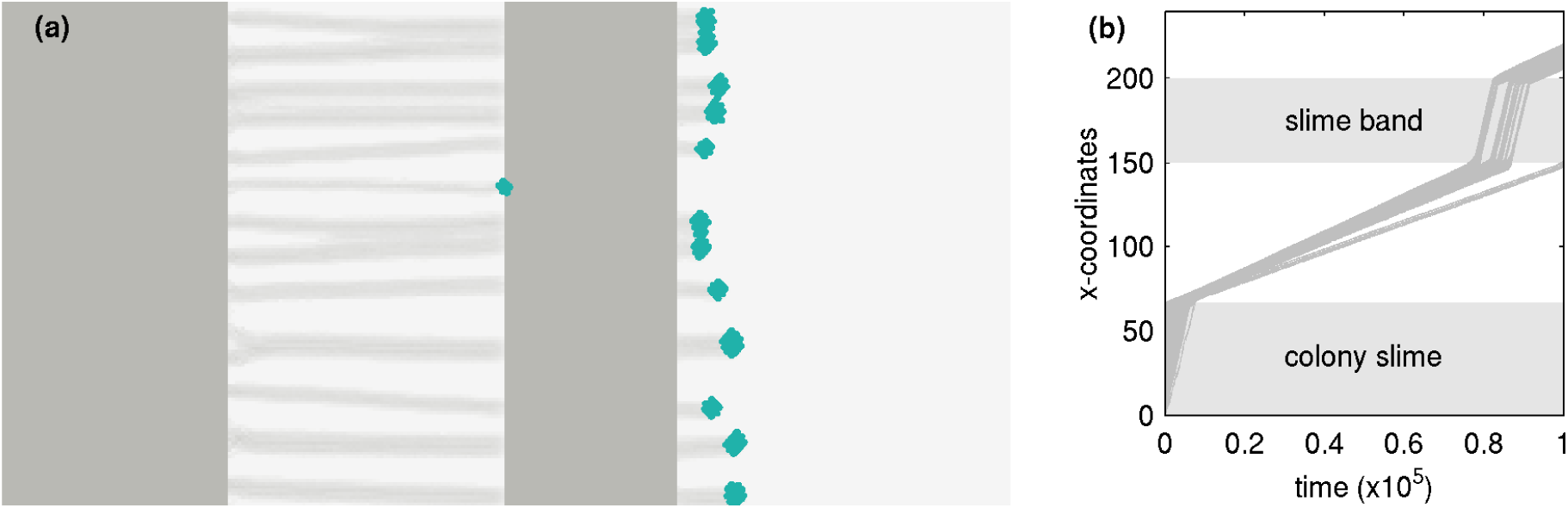
Movement of fingers through a slime band: (a) Motion of fingers through a slime band after extending from a flat colony. (b) Kymograph of cells indicating how velocities increase when cells in fingers encounter a pre-existing band of slime.

Fig. 6 shows the evolution of fingers from an initial flat colony that contains a fraction of “freeloaders” mixed with ordinary cells. These freeloaders do not sense light although, they can move and lay down slime. Their net motion towards light can thus only come from being dragged along by cells that do sense light. Our results show that these freeloaders can in fact entrain with cells that sense and move towards light, but the precise details of finger formation and evolution depend both on *p*_photo_ and the initial concentration of freeloaders *ϕ*_ch_. At small *p*_photo_ (Fig. 6 (a,d,g)) one finds small and disordered fingers, whereas at intermediate values of *p*_photo_ = 0.10 (Fig. 6(b,e,h)), well developed fingers are observed even at somewhat larger *ϕ*_ch_. If *p*_photo_ is large (Fig. 6(c,f,i)), entrainment is less successful and we find that the freeloaders can be left behind in the colony, even as normal cells move toward light in fingers. When the tip of a finger comprises both types of cells, we observe that freeloaders tend to cluster towards the back of such tips.

**Figure 6:**
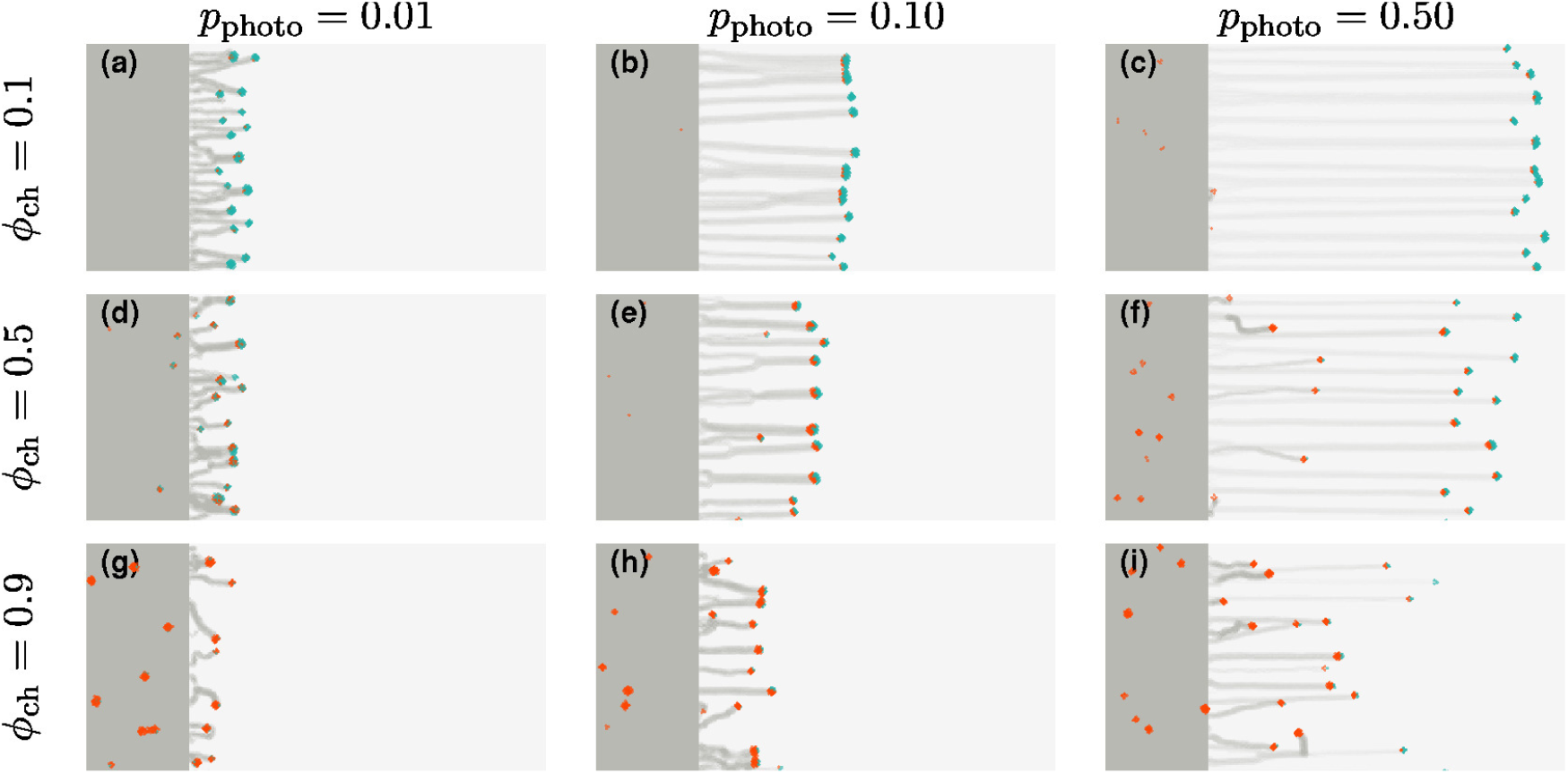
“Freeloaders” that do not sense light can also move toward light through entrainment: Snapshots of the colonies at *t* = 10^5^ for a range of *p*_photo_ and for different ratios of cells that do not sense the direction of light (freeloaders, at a fraction *ϕ*_ch_ of the total number of cells, represented in red). We consider the cases (a-c) *ϕ*_ch_ = 0.1, (d-f) *ϕ*_ch_ = 0.5, and (g-i) *ϕ*_ch_ = 0.9. For each of these values of *ϕ*_ch_, we show snapshots for (a, d, g) *p*_photo_ = 0.01, (b, e, h) *p*_photo_ = 0.1 and (c, f, i) *p*_photo_ = 0.5. Fingers can be observed even for high values of *ϕ*_ch_, especially for high values of *p*_photo_. We observe that freeloaders cluster towards the back of the tips of fingers.

The advantage of a simple model is that it provides predictions for more complex cases in which the position of the light source changes over the course of the experiment. We are able to replicate observations of fingers that turn when the source of light is moved [26]. In Fig. 7(a), we show how fingers extend from the colony towards the direction of a source of light, initially placed towards the right of the colony. After these fingers have developed substantially, the position of the light source is changed instantaneously so that light now emanates from a point rotated 90^*?*^ clockwise from the original position. As can be seen in Fig. 7(b), there is a sharp kink that develops in the fingers as the cells at the tip change their direction of motion so as to follow the light. The fingers now develop and extend in the new direction of light (See Movie S3).

**Figure 7:**
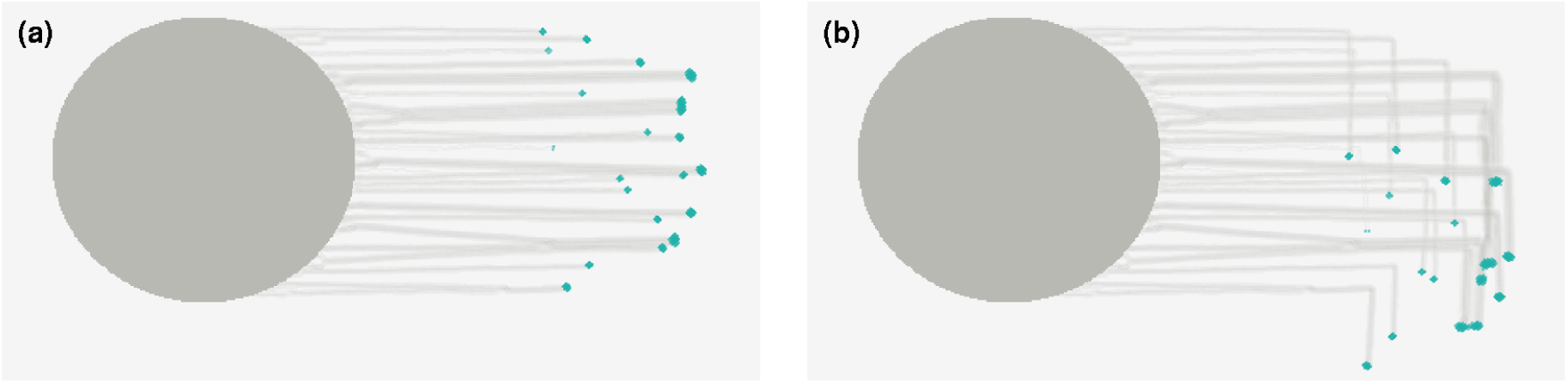
Changing the direction of light: Cells in a colony of density *ρ* = 0.2 move toward a light source to the right until *t* = 2 ×10^5^ at which time the position of light source is moved 90 *°* clockwise. The cells reorient themselves and start moving toward the new direction of light. (a) Snapshot at *t* = 2 × 10^5^ (b) Snapshot at *t* = 2.5 × 10^5^.

## Discussion

Our agent-based model for the phototaxis of *Synechocystis* reproduces prior observations concerning the finger-like projections that form and extend in colonies of motile cells as a phototactic response. The fundamental unit in our model is a single cell which interacts with its environment by sensing light. Two central features of our model are the ability of motile cells to lay down slime, thereby facilitating the motion of cells that subsequently traverse that region, as well as the ability of T4P to mediate the direct physical interaction of cells. These interactions lead to collective behaviour in the form of aggregation and subsequent finger formation, as cells migrate in the direction of light. We suggest that this interaction is central to collective phototaxis.

Cell-cell interactions and the ability of cells to self-propel are essential aspects of active matter descriptions of collective cell migration. Our model can be viewed as an extension of studies in which net motion depends on an externally imposed cue, rather than arising from a spontaneous breaking of symmetry. The presence of slime is an ingredient of our model, which appears to have no direct precedent in active matter models. (A possible exception are models for ant trails, which rely on signalling through pheromones laid down by ants that sample multiple paths [40].) The advantage of these and similar mathematical and computational models is that, once benchmarked, they allow us to query aspects of experimental systems that can also be separately examined in targeted experiments(as in [41]). In addition, they allow us to estimate ranges of e.g., force parameters (*k*_1_*, k*_2_) and the slime deposition rate (*S*_rate_) that can generate fingers reminiscent of ones observed in experiments. Interestingly, relatively small values of *p*_photo_ appear to be sufficient to generate well-developed fingers; a result that is in agreement with experimental observations [42].

Chemotaxis-like gene clusters in bacteria such as *Synechocystis* have several photoreceptors [12, 43, 44]. These clusters also contain genes involved in signal transduction, including genes that code for signal receptors, response regulators and motility regulators [44, 45]. A number of motility-related genes involved in the biosynthesis and function of pili are required for phototaxis [36, 37, 46]. Genes involved in cAMP regulation have also been shown to be important in phototaxis [47]. Specifically, phototaxis was found to be impaired in colonies of mutants in *cya1* (an adenylate cyclase) and *sycrp1* (a cAMP receptor-like protein), which accumulate near the colony edge closest to a light source but do not extrude outwards in fingers [18]. We model such mutants as “freeloaders”, since they cannot, on their own, exhibit the directional motion characteristic of phototaxis. However, provided the cell-cell interactions are intact, one can expect that combining a small density of freeloaders with other cells capable of sustained phototaxis might be sufficient to sustain motion of the collective. We have shown that the role of freeloaders, as well as the consequences of a change in light direction, can be examined systematically.

Other scenarios can be tested in models and then later investigated through experiments. These include effects of multiple light sources and wavelengths, as well as finger formation in mutants that cannot either move or produce slime. Whether such taxis mutants might lead to different architectures of colonies remains to be fully explored. In principle, expanding this model further to include a quantification of fitness might also allow us to address evolutionary questions, such as whether a fraction of freeloaders can persist in mixed populations over several generations. Finally, we note that the term quorum-sensing is conventionally applied to a situation where a commonly sensed biochemical signal crosses an activation threshold. Thus, quorum-sensing is at its core a collective process. Phototaxis as discussed here also embodies a collective effect in the TFP-mediated cell-to-cell interaction as well as slime-mediated interactions. For this reason it may be worth exploring other contexts for quorum-sensing that emphasize its origins in collective, especially physical interactions, and nonlinear response.

In conclusion, we have developed a model for phototaxis in *Synechocystis* colonies which incorporates several features that underlie collective motion in systems of this nature. Our model describes the movement of individual cells, each having a finite volume and capable of detecting light. Cell motion is accompanied by the deposition of slime that serves to reduces friction. The role of T4P is captured in our model by allowing cells to attach to neighbours, and exert forces on them. We observe that the collective behaviour is characterized by cells aggregating into small clusters that first accumulate at the edge of the colony. These clusters then extrude towards the light source in finger-like projections, reminiscent of recent experimental observations. We find that these projections occur even when the bias of individual cell motion towards light is very small, and also in situations where some fraction of the colony consists of freeloaders that do not sense the direction of the light source. Further improvements of our model would involve the coupling of the systems biology of light-sensing with our physical model for cyanobacterial motility.

## Acknowledgements

We would like to thank Sandeep Krishna, Rajesh Singh and Shashi Thutupalli for helpful discussions. S.N.M. is supported by the IMSc Complex Systems Project (12^th^ Plan). We thank IMSc for providing access to the supercomputing cluster “Satpura”, which is partially funded by DST.

